# A comparison of *Dictyostelium discoideum* 3’-5’ RNA polymerases reveals a conserved tRNA^His^ guanylyltransferase residue that plays a dual role in catalysis

**DOI:** 10.1101/2025.10.12.681950

**Authors:** Grace N. Johnecheck, Korey J. Kihm, Jane E. Jackman

## Abstract

The 3’-5’ RNA polymerase family consists of eukaryotic tRNA^His^ guanylyltransferase (Thg1) and Thg1 homologs known as Thg1-like proteins (TLPs) that exist in all three domains of life. Thg1 catalyzes an essential reaction adding a G-1 nucleotide to the 5’ end of the tRNA^His^, forming an identity element for tRNA aminoacylation. All TLPs studied, except *Dictyostelium discoideum* (Ddi) TLP2, perform *in vitro* Watson-Crick (WC) dependent addition of multiple nucleotides to repair truncated tRNA. DdiTLP2 has a similar activity to a Thg1 enzyme, adding G-1 to mt-tRNA^His^, but shares other biochemical properties with other TLPs, including a restriction to making WC base pairs during this reaction. We identified two regions in DdiTLP2 that lacked residues that are absolutely conserved in other Thg1/TLP enzymes. DdiTLP2 variants in both regions abolish enzymatic activity of DdiTLP2, indicating these regions are important for DdiTLP2 catalysis. Complementary alterations to one of these residues (D150) in DdiThg1 caused an unexpected reversal of this enzyme’s specificity, with a loss of its ability to incorporate a non-WC base paired G-1 to its physiological substrate, while gaining the ability to add WC base paired G-1 to mt-tRNA^His^. These biochemical results, combined with structural models suggest a previously unknown role for D150 in controlling substrate specificity at the adenylation step by providing a checkpoint for correct setup of a WC base pair in the active site. Thg1 also appears to have adapted the role of the ancestral D150 residue for a second function, promoting non-WC nucleotide addition to its eukaryotic tRNA^His^ substrate.

## INTRODUCTION

Unlike most polymerases that act in the 5’ to 3’ direction, members of the tRNA^His^ guanylyltransferase (Thg1) enzyme family synthesize RNA 3’ to 5’. The founding member of the Thg1 family in *Saccharomyces cerevisiae* (ScThg1) is an enzyme that adds a single G nucleotide to the 5’ end of tRNA^His^ (Gu et al. 2003; Jackman and Phizicky 2006a). In most eukaryotes, this essential G nucleotide (known as G-1 due to its location prior to the typical 5’- end of all other tRNAs) acts as the identity element for histidyl-tRNA synthetase (HisRS) to recognize tRNA^His^ and charge it with histidine (Cooley et al. 1982; Rudinger et al. 1994; Rudinger et al. 1997). The essential function of Thg1 in eukaryotes is to perform template-independent 3’ to 5’ addition of this critical G-1 nucleotide across from the tRNA^His^ A_73_ nucleotide, thus forming a non-Watson-Crick (WC) G-1:A73 base pair at the top of the aminoacyl acceptor stem of tRNA^His^ **(Figure 1A)** (Gu et al. 2005; Preston and Phizicky 2010).

**Figure 1.**
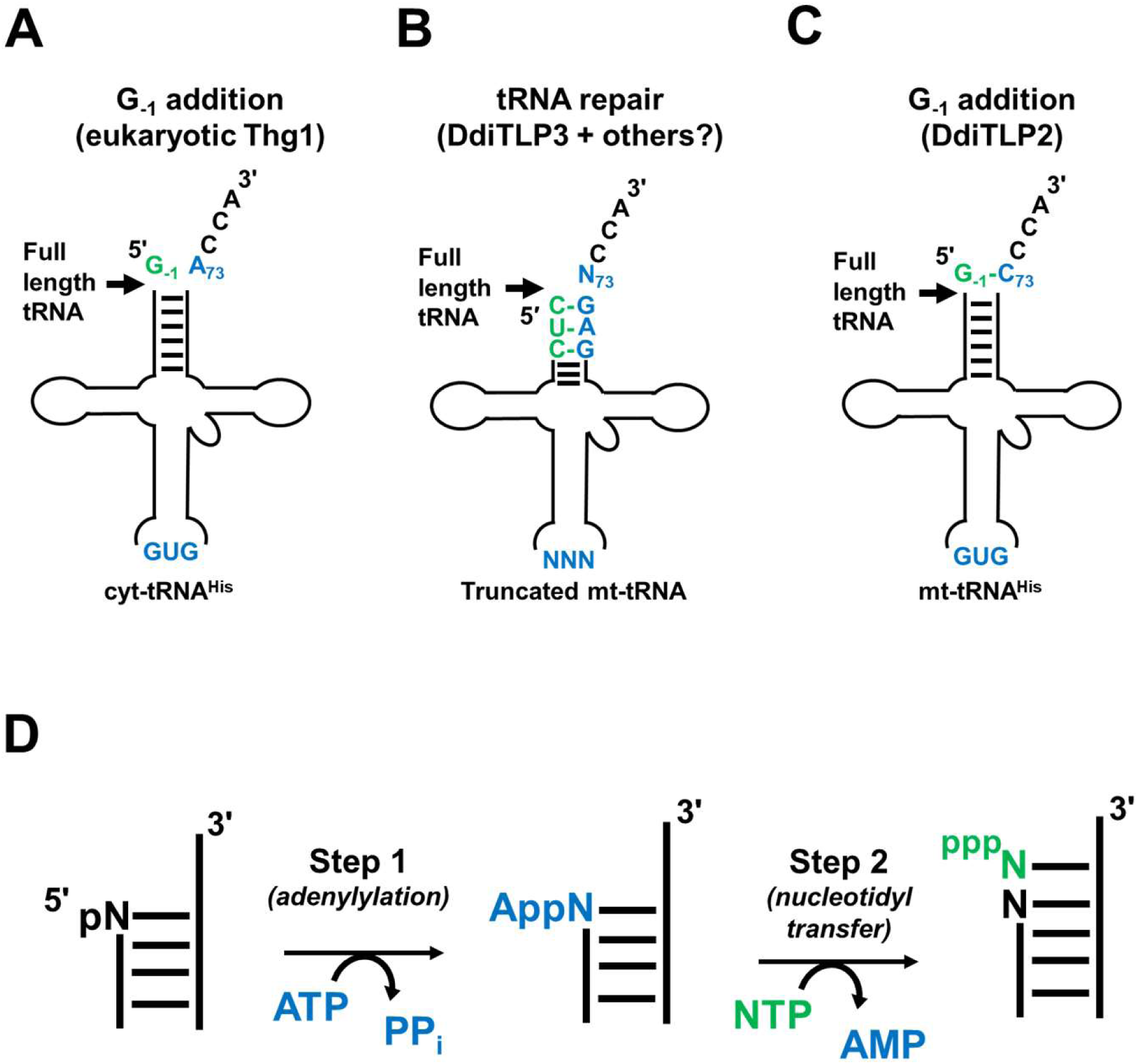
Three known biological functions for Thg1/TLP 3’-5’ RNA polymerases. For each indicated tRNA, black arrows indicate the size of full-length mature tRNA species, which is relevant to the site of addition for each type of enzyme. Specific tRNA sequences that are relevant for each indicated activity are shown in blue. Nucleotides incorporated by Thg1/TLP enzymes during each reaction are indicated in green. **A) G_-1_ addition by eukaryotic Thg1.** Thg1 adds a single G-1 nucleotide to the 5’ end of cyt-tRNA^His^, extending the full-length tRNA by one nucleotide. Substrate recognition requires the presence of the His GUG anticodon, and G-1 is incorporated in a non-WC base pair across from the A73 discriminator nucleotide. **B) tRNA repair by DdiTLP3.** DdiTLP3 adds nucleotides to 5’-truncated mt-tRNAs that are generated by the removal of mismatched nucleotides from the 5’-end of the tRNA during 5’-editing. The number of nucleotides added during repair can vary for different tRNAs, and results in a full-length tRNA with a completed 7 base pair aminoacyl-acceptor stem. This activity does not require any specific anticodon sequence and therefore occurs on multiple tRNAs of different types in species where it has been characterized. Nucleotides added by DdiTLP3, and all other TLPs that can catalyze this same activity *in vitro*, are incorporated in a WC base pair dependent manner, with the nucleotides on the 3’-half of the tRNA (GAG shown in blue here) acting as a template for the 3’-5’ RNA polymerase activity. **C) G_-1_ addition by DdiTLP2.** DdiTLP2 adds a single G-1 nucleotide to the 5’ end of mt-tRNA^His^, also resulting in extension of the full-length RNA by one nucleotide. Substrate recognition by DdiTLP2 does not require the GUG anticodon, and instead relies on other identity elements that remain to be identified. G-1 is incorporated in a WC template-dependent manner, forming a G-1-C73 base pair after addition. **D) Generalized two-step 3’-5’ nucleotide addition reaction.** In step 1 of the reaction with a 5’-monphosphorylated RNA substrate, Thg1/TLP enzymes adenylylate the 5’-end using ATP. In step 2, Thg1/TLP enzymes transfer the indicated NTP to the 5’-end, using the 3’-OH of the incoming NTP for nucleophilic attack on the activated intermediate, thus resulting in the 5’-extended RNA. Because the 5’-triphosphate of the added NTP is retained during step 2, addition of subsequent NTPs during multiple nucleotide addition does not require re-activation of the substrate. Instead, the 5’-triphosphate provides the necessary leaving group for nucleophilic attack by the incoming NTP, releasing pyrophosphate and generating a new triphosphorylated end for the subsequent NTP addition.

Thg1 is part of a larger family of 3’ to 5’ polymerases found in all three domains of life (Jackman and Phizicky 2006b; Abad et al. 2010; Chen et al. 2019). ScThg1 was the first enzyme shown to be capable of using 3’-5’ addition activity to create WC base-paired nucleotides, like a canonical 5’-3’ polymerase, in an *in vitro* reaction that is distinct from its required non-WC G-1 addition activity (Jackman and Phizicky 2006b). Subsequent biochemical studies revealed that templated 3’-5’ polymerase activity is a likely ancestral feature shared by all members of the enzyme family (Abad et al. 2010; Rao et al. 2011; Desai et al. 2018; Dodbele et al. 2019). Many enzymes have now been characterized that comprise a distinct group of proteins with representatives in Eukarya, Archaea and Bacteria, known as Thg1-like proteins (TLPs) (Abad et al. 2011; Jackman et al. 2012; Chen et al. 2019). TLPs lack the eukaryote-specific activity of Thg1 to add the non-WC base-paired G-1 to tRNA^His^ (**Figure 1A**) and instead TLPs from all domains of life only exhibit WC-dependent 3’-5’ addition activity *in vitro* and *in vivo* (**Figure 1B** and **1C**) (Abad et al. 2010; Rao et al. 2011; Desai et al. 2018). Thus, the ability of Thg1 homologs to catalyze two distinct types of 3’-5’ addition activities (WC and non-WC) appears to be a derived trait in the eukaryotic lineage.

One reaction demonstrated for TLPs involves the use of multiple 3’-5’ additions to repair 5’-truncated tRNAs that are missing nucleotides from their 5’-end **(Figure 1B)**. Like the templated reactions catalyzed by canonical DNA or RNA polymerases, TLPs use the 3’-end of the tRNA as a template to select the correct WC base-pairing NTP to be added to the 5’ end of the tRNA, in this case restoring a complete 7 bp aminoacyl acceptor stem(Abad et al. 2011; Betat et al. 2014; Long et al. 2016). In the slime mold *Dictyostelium discoideum*, at least one TLP uses this tRNA 5’-end repair activity physiologically to repair mitochondrial-encoded tRNA (mt-tRNA), as part of a process that is known as tRNA 5’-editing (Abad et al. 2014; Long et al. 2016). Other protozoan eukaryotes that encode TLPs require a similar mt-tRNA 5’-editing activity, and therefore TLP orthologs in these species predictably also utilize the 3’-5’ polymerase reaction for this function (Lonergan and Gray 1993a; Lonergan and Gray 1993b; Burger et al. 1995; Laforest et al. 1997; Laforest et al. 2004; Bullerwell and Gray 2005).

The first *in vivo* characterization of any TLP enzymes was in the eukaryotic slime mold *D. discoideum*, which encodes four different Thg1/TLP enzymes, three of which are essential for viability (Long et al. 2016). In *D. discoideum*, one of these essential enzymes is a single Thg1 ortholog (DdiThg1), which adds G_-1_ to cytosolic (cyt-) tRNA^His^, as observed in other eukaryotes (**Figure 1A**). The other three genes in *D. discoideum* encode TLP-type enzymes (DdiTLP2, DdiTLP3 and DdiTLP4). DdiTLP3 comprises part of the mt-tRNA 5’-editing machinery, where its essential function is to use templated 3’-5’ polymerase activity to repair multiple 5’-truncated mt-tRNAs that are generated by removal of incorrectly-encoded mismatched nucleotides from the 5’-end of the precursor mt-tRNA (**Figure 1B**) (Abad et al. 2011; Abad et al. 2014; Long et al. 2016). Interestingly, DdiTLP4 is also an essential enzyme in *D. discoideum* and catalyzes a similar biochemical tRNA repair reaction to DdiTLP3 *in vitro*, yet DdiTLP4 is present in the cytosol where there are not known to be any truncated mt-tRNA that could be substrates for this kind of activity. Thus, the essential function of DdiTLP4 in *D. discoideum*, as well as the function of many other TLPs in organisms that also lack predictable sources of truncated tRNA, remains unknown.

The unique nature of the third *D. discoideum* TLP, DdiTLP2, was suggested by the observations that 1) DdiTLP2 is not an essential enzyme in *D. discoideum* (although its deletion causes a slow growth phenotype) and 2) that it cannot repair truncated tRNA *in vitro* like all other TLP-type enzymes characterized to date (Long et al. 2016). Instead, the demonstrated physiological role of DdiTLP2 is to add a G-1 residue selectively to mt-tRNA^His^, which makes its activity more similar to that of Thg1-type enzymes (**Figure 1C**). However, despite these similarities to Thg1, DdiTLP2 also possesses several biochemical characteristics that reflect its classification into the TLP-type group. First, like all TLPs studied to date, DdiTLP2 strongly prefers to create a WC base pair during G-1 addition (**Figure 1C**), and lacks the characteristic non-WC addition activity of Thg1 orthologs (**Figure 1A**) (Long et al. 2016). Second, although Thg1-type enzymes so far all depend on the tRNA^His^ GUG anticodon for recognition of their cognate tRNA (Jackman and Phizicky 2006a; Jackman and Phizicky 2008), we observed that DdiTLP2 does not require the GUG sequence to add G-1 to its substrate mt-tRNA^His^ (Long et al. 2016). Thus, like canonical TLPs, DdiTLP2 does not require a specific anticodon for its activity and therefore must utilize a distinct strategy to selectively recognize its cognate mt-tRNA^His^.

The molecular basis for these unique features of DdiTLP2 that blend Thg1-like properties (G-1 addition to full-length tRNA) and TLP-like properties (WC base pair dependence and lack of recognition of GUG) are not known. Moreover, although isolated amino acids that impact the biochemical properties of Thg1 have been identified (Jackman and Phizicky 2008; Smith and Jackman 2012; Nakamura et al. 2013; Nakamura et al. 2018; Matlock et al. 2019; Patel et al. 2021), a complete understanding of the features that dictate the distinct biochemical behaviors of Thg1- vs. TLP-type enzymes overall has been elusive. Despite the availability of several structures of both types of enzymes, including two complexes with RNA, many questions about the molecular basis of base pair recognition and tRNA specificity remain unanswered (Hyde et al. 2010; Hyde et al. 2013; Nakamura et al. 2013; Kimura et al. 2016; Nakamura et al. 2018). As Thg1/TLPs are the only known enzymes capable of performing templated nucleotide addition to the 5’-ends of RNA substrates, obtaining a complete picture of the mechanistic basis for this activity will facilitate future approaches to potentially exploit this activity. Thus, the *D. discoideum* enzymes provide us with an excellent system to try to address these questions, since there are three distinct patterns of biochemical activity represented among these family members.

Here we used sequence comparison and biochemical approaches to test functions for specific residues DdiTLP2 vs. other Thg1/TLP enzymes. We identified two unique regions of DdiTLP2 and created variants to test the functions of these alterations on the enzymatic properties of the resulting recombinant proteins. One stretch of residues in a conserved loop structure appears to be critical for DdiTLP2’s catalytic function. We also investigated an unusual single amino acid change in DdiTLP2 involving an aspartate that is otherwise universally conserved as aspartate or glutamate among Thg1 and TLP family members. Biochemical and kinetic characterization of the conserved D150 revealed that this residue plays a dual role in DdiThg1, acting both as a gatekeeper to ensure fidelity of 3’-5’ addition activity and to specifically promote the specialized function of DdiThg1 in G-1 addition to tRNA^His^. These two functions exploit distinct chemical properties of the aspartate side chain. A combination of structural models with the biochemical results obtained here suggest a previously unknown function of this D150 residue in the coupled two step reaction mechanism that was recently elucidated for Thg1/TLP enzymes, and help to answer long-standing questions about the evolution of eukaryotic Thg1 activity and its impact on biology.

## RESULTS AND DISCUSSION

### DdiTLP2 contains unique residues that distinguish it from other 3’-5’ RNA polymerases

The biochemical properties of DdiTLP2 suggest that it occupies a unique position in the 3’-5’ RNA polymerase family (**Figure 1**). Thus, we sought to identify sequence features that are unique to DdiTLP2 and that could distinguish this enzyme from its canonical Thg1 and/or TLP orthologs. Sequence alignment was performed between DdiTLP2 and eleven other Thg1/TLP enzymes from all three domains of life whose biochemical activities have been characterized (Abad et al. 2010; Heinemann et al. 2010; Hyde et al. 2010; Abad et al. 2011; Smith and Jackman 2012; Hyde et al. 2013; Nakamura et al. 2013; Rao et al. 2013). These enzymes included four canonical “Thg1”-type enzymes that add G-1 to nucleus-encoded tRNA^His^ (**Figure 1A**): *Homo sapiens* (Hs) Thg1, *Saccharomyces cerevisiae* (Sc) Thg1, *Candida albicans* (Ca) Thg1 and *Dictyostelium discoideum* (Ddi) Thg1, and seven “TLP”-type enzymes that all, unlike DdiTLP2, repair 5’-end truncated tRNA *in vitro* (**Figure 1B**): *D. discoideum* TLP3 and TLP4, *Acanthamoeba castellanii* (Aca) TLP1 and TLP2, *Bacillus thuringiensis* (Bt) TLP, *Methanosarcina acetivorans* (Ma) TLP; and *Methanopyrus kandleri* (Mk) TLP **(Figure 2 and Figure S1).**

**Figure 2.**
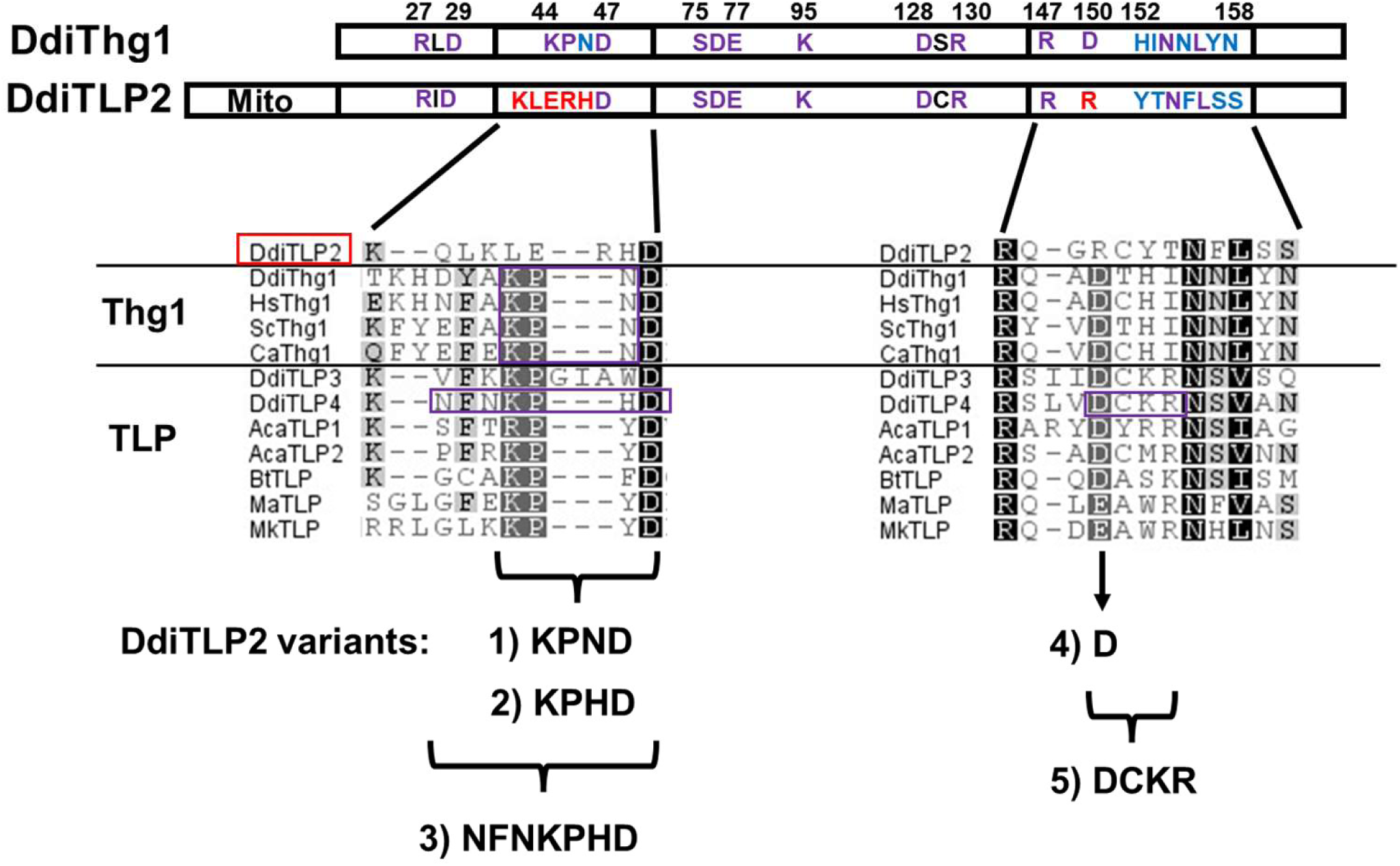
Thg1/TLP multiple sequence alignment reveals unique DdiTLP2 sequences in regions of otherwise strong Thg1/TLP conservation. A comparison of DdiThg1 and DdiTLP2 shows key residues conserved in both Thg1 and TLP enzymes indicated in purple, residues uniquely conserved in Thg1 indicated in blue, and residues unique to DdiTLP2 indicated in red (See **Figure S1** for complete sequence alignments). Residues numbering is according to locations in the DdiThg1 sequence. The region labeled “Mito” indicates a predicted N-terminal mitochondrial transport sequence in DdiTLP2. Detailed alignments showing two regions of the Thg1/TLP where DdiTLP2 diverges from other family members are shown below each highlighted sequence. The sequence alignment was performed as described in Methods for Thg1/TLP enzymes from all three domains of life: *Dictyostelium discoideum* (Ddi) Thg1, TLP2, TLP3, and TLP4; *Bacillus thuringiensis* (Bt) TLP; *Acanthamoeba castellanii* (Aca) TLP1 and TLP2; *Homo sapiens* (Hs) Thg1; *Saccharomyces cerevisiae* (Sc) Thg1; *Candida albicans* (Ca) Thg1; *Methanosarcina acetivorans* (Ma) TLP; and *Methanopyrus kandleri* (Mk) TLP. The five DdiTLP2 variants created for this work are indicated by brackets with each corresponding replacement sequence.

As expected, given the core 3’-5’ RNA polymerase activity exhibited by all twelve enzymes, there are many regions of significant conservation across all enzymes, including key residues whose function in Thg1/TLP catalysis has been established (**Figure 2**, **Figure S1**).

These include the metal ion binding carboxylates (DdiThg1 D29, D76, and E77; residue numbering used in this manuscript is for the corresponding amino acids in DdiThg1, unless otherwise indicated), residues that are required for efficient 5’-adenylylation (K44 and S75), and many residues whose precise molecular function has not been established but that are known play critical roles in catalysis for various enzymes in the family (R27, K95, D128, R130, R147). Despite their strong conservation, alanine variants of several amino acids (D47 and N154) in the context of ScThg1 do not disrupt catalytic activity, either *in vitro* or *in vivo* in *S. cerevisiae* (Jackman and Phizicky 2008). Nonetheless, roles for these residues in other enzymes or other Thg1 functions may yet be demonstrated. Finally, several sequences are known to confer the separate, non-WC, 3’-5’ addition activity onto eukaryotic Thg1 enzymes, and thus share strong conservation between Thg1 enzymes but are not found in the *bona fide* repair TLPs. For example, the Thg1-specific HINNLYN sequence contains 3 catalytically important residues, including H152, Y157 and N158 residues (Jackman and Phizicky 2008; Smith and Jackman 2012). These amino acids are essential for Thg1’s physiological non-WC base paired addition of GTP across from A73 in tRNA^His^ (**Figure 1A**), but are not required for its ability to catalyze WC base paired addition activity at the same position (Matlock et al. 2019; Patel et al. 2021). These data suggest that acquisition of these eukaryotic Thg1-specific residues was an important part of the specialization of these enzymes for their known function in tRNA^His^ maturation.

We identified two regions where the DdiTLP2 sequence differed significantly from generally well-conserved sequences in both Thg1 and TLPs, and we hypothesized these could be residues associated with the unique biochemical properties of DdiTLP2. The first region is in a loop that contains the otherwise highly conserved KPXD sequence, in which K44, P45 and D47 are conserved in all other Thg1/TLP enzymes (**Figure 2, Figure S1**). In the crystal structures of all Thg1/TLPs to date, the KPXD sequence is part of a short (6-8 amino acid) loop that connects two alpha helices near the active site. AlphaFold3 models of DdiTLP2, DdiThg1 and DdiTLP4 predict a similar loop structure for each enzyme, with each loop ending at the conserved D47 (**Figure 3**). The KPXD-containing loop surrounds the ATP nucleotide that is used for step 1 of the Thg1/TLP reaction (**Figure 1D**), consistent with the observed kinetic role for K44 in 5’-adenylylation. However, none of the loop residues are observed in these models making consistent interactions with the ATP that would provide insight into their actual molecular role(s), which also remain unexplained by existing structures that contain bound ATP at the adenylylation site (Hyde et al. 2010; Hyde et al. 2013; Kimura et al. 2016). The X residue is always N46 in Thg1 enzymes, while in TLPs, X46 is more variable, but is often an amino acid with some aromatic character (Y,F,W,H). An ScThg1 N46Y variant retains its ancestral ability to polymerize WC base pairs *in vitro*, while losing its ability to catalyze the Thg1-specific non-WC G-1 activity (**Figure 1A**), consistent with the idea that the acquisition of N46 in the eukaryotic lineage helped enable evolution of the non-templated G-1 addition reaction catalyzed exclusively by Thg1 orthologs (Patel et al. 2021). Interestingly, DdiTLP2 deviates substantially from both Thg1 and TLPs in this loop region, lacking residues that correspond with K44 or P45, and retaining only the conserved D47 and a possible TLP-like H46 (**Figures 2 and 3**).

**Figure 3.**
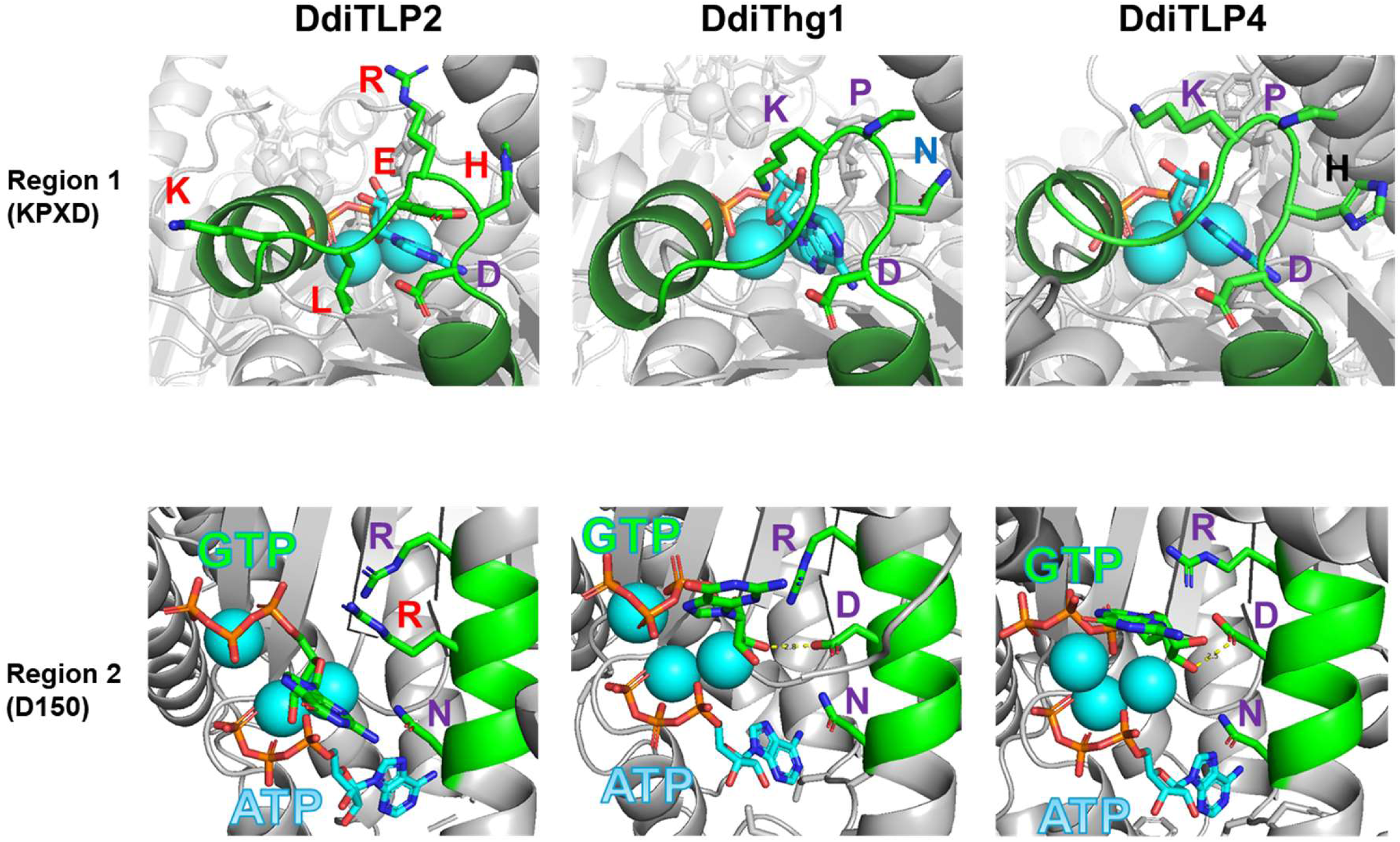
AlphaFold3 models of unique regions identified from sequence alignments. DdiTLP2, DdiThg1 and DdiTLP4 were each modeled using AlphaFold3 with two tRNAs, plus three metal ions (cyan), one GTP (green) and one ATP (cyan) per monomer, consistent with the stoichiometries observed in structures of enzymes obtained to date (see Methods for details). The tRNA has been removed from the views shown here for each region for clarity. Residues are colored according to the same scheme used in Figure 2 (purple, conserved in all enzymes; red, unique to DditLP2, blue, unique to DdiThg1). Region 1 shows the KPXD-containing loop region that surrounds the ATP bound in the 5’-adenylylation (step 1) site, which in all models terminates with the universally conserved D47. Despite its apparent structural conservation, D47 is not essential for ScThg1 activity, *in vitro* or *in vivo* (Jackman and Phizicky 2008). Region 2 shows a segment of alpha helix that contains universally conserved R147 and N154 pointed into the active site towards an NTP that has been identified kinetically as corresponding to the incoming GTP for nucleotidyl transfer (step 2). Potential hydrogen bonding interaction between D150 and the 3’ OH of GTP is shown by a dotted yellow line (2.8 Å for DdiThg1 and 2.5 Å for DdiTLP4). Full models of each enzyme-tRNA complex are shown in **Figure S2**.

In the second region, a conserved D/E residue (D150 in DdiThg1) is invariant among all studied Thg1/TLP enzymes aside from DdiTLP2, which contains an R187 at this location (**Figure 2 and 3**). Although structures of DdiTLP2 have not been experimentally determined, the AlphaFold models of DdiTLP2 all show this R (R187 in DdiTLP2) adopting an analogous position to D150 within a highly conserved alpha helix that positions three nearly invariant residues (R147, D150, N154) all pointing toward the GTP nucleotide that will be transferred to the 5’-end during the second, nucleotidyl transfer, step of the reaction (**Figure 3**, **Figure 1D**).

This alpha helix with similarly arranged amino acids has been observed in every structure of a Thg1 or TLP solved to date, yet the specific roles of these conserved amino acids have not been evident from any of the captured complexes. Interestingly, the analogous D150A variant of ScThg1 (D153 in ScThg1) has been tested previously. This variant exhibited a puzzling phenotype, with catalytic (G-1 addition) activity *in vitro* that was actually higher than the wild-type enzyme, while yeast cells that express only the D153A variant version of this essential enzyme exhibited an obvious growth defect characterized by slow growth and variable colony size. No subsequent structural or biochemical studies have provided any further insight into the molecular role of D150. Thus, we sought to investigate whether the distinct sequence patterns in these two motifs could be involved in DdiTLP2’s unique enzymatic functions.

### DdiTLP2-specific residues are important for catalytic activity on mt-tRNA^His^

To test the relevance of these unusual amino acids for DdiTLP2 activity, we created variants to exchange sequences to those found in either Thg1 or other TLP orthologs (**Figure 2**). We created two variants of DdiTLP2 in Region 1 in which the KLERHD sequence was replaced by KPND, corresponding to Thg1, or KPHD, corresponding to DdiTLP4. A third variant was also created with a longer part of the sequence (QLKLERHD) was replaced with the entire loop sequence predicted for DdiTLP4 (NFNKPHD), to allow for variation in the loop length. Two additional alterations were made in Region 2 of DdiTLP2. First, the R187D variant was created, restoring the otherwise universal D150 residue to this alpha helix. Second, a four amino acid change was made in the second half of the helix to match the sequence of this entire helical turn to that found in both DdiTLP3 and DdiTLP4 (RCYT to DCKR). Based on AlphaFold3 structural predictions, these changes would predictably maintain the helical structure with the conserved D150 pointed toward the modeled GTP nucleotide that is in the nucleotidyl transfer site, as in all other Thg1/TLP enzymes solved or predicted to date.

All five variant enzymes were expressed in and purified from *E. coli*, and nearly all of the enzymes were purified with relatively high yield, except for the longer loop variant (Variant 3), which may indicate issues with protein folding **(Figure S2)**. The variants were then tested for their activity with different 5’-^32^P-end labeled tRNA substrates using a phosphatase protection assay. In this assay, addition of 5’-nucleotide(s) to the substrate RNA creates a phosphatase-resistant oligonucleotide product after nuclease digestion (see methods). Products of 3’-5’ nucleotide addition can then be resolved from inorganic phosphate (derived from unreacted substrate) using thin layer chromatography. Assays were first performed with *D. discoideum* mitochondrial tRNA^His^ (mt-tRNA^His^), where the labeled fragment GpGpU corresponds to the product of G-1 addition. As demonstrated previously, DdiTLP2 exhibits robust in vitro activity with this tRNA, while wild-type DdiThg1 activity is weak, consistent with each enzyme’s known biological function (**Figure 4A**) (Long et al. 2016). All five of the changes made to DdiTLP2 completely disrupted its catalytic activity with the mt-tRNA^His^ substrate, with only a very faint amount of product visible in the assays with the highest concentration of the KPHD variant (**Figure 4A**).

**Figure 4.**
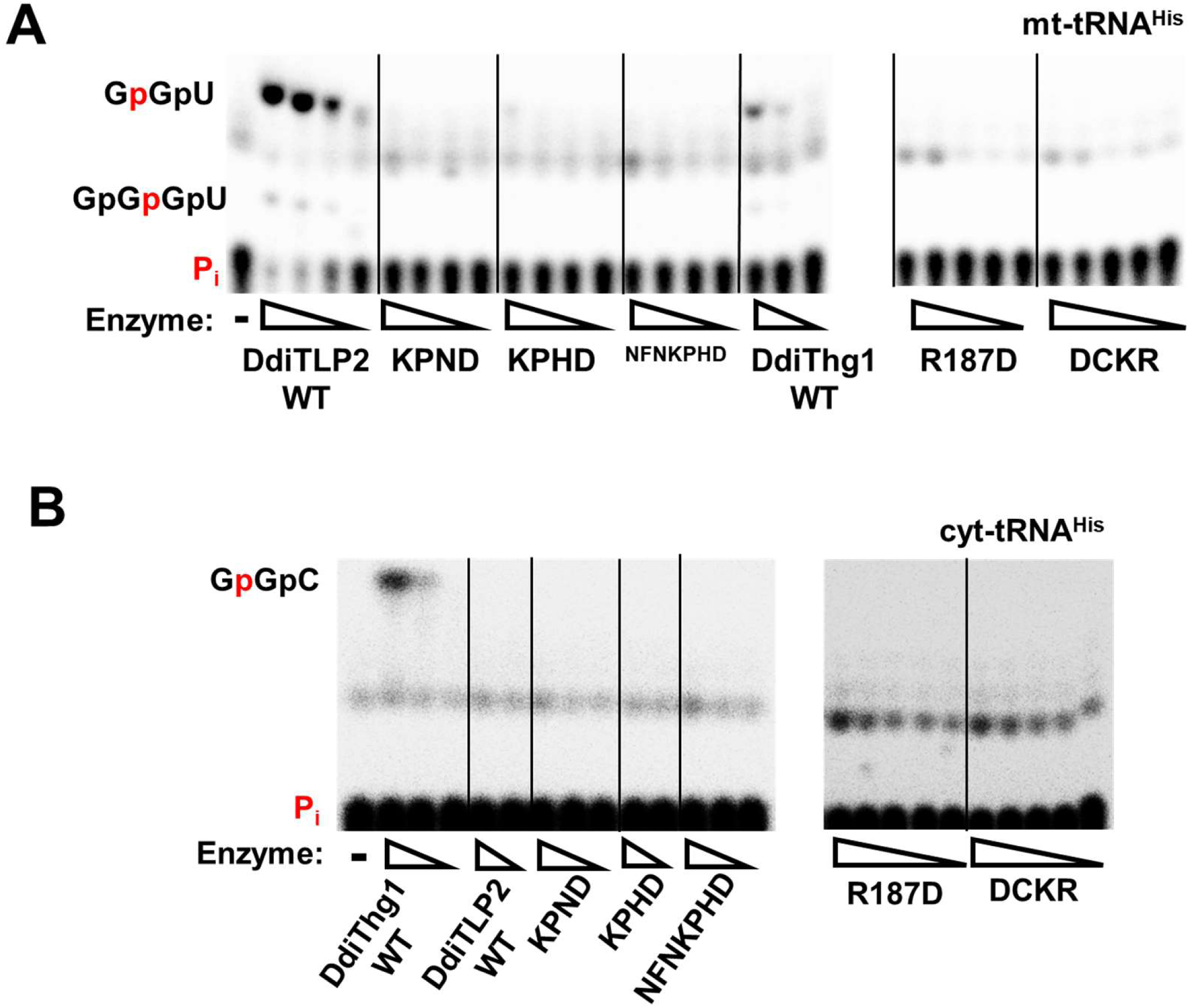
Residues that are unique to DdiTLP2 are required for catalytic activity. Phosphatase protection assays were performed with **(A) Ddi mt-tRNA^His^** and **(B) Ddi cyt-tRNA^His^** using 5-fold serial dilutions of the indicated enzymes. Reactions contained 0.1 mM ATP, 1 mM GTP, and 5’-^32^P-labeled tRNA and were quenched after 2 hour incubation by addition of EDTA, followed by RNase A and phosphatase treatment to digest any reaction products into shorter fragments, as shown on the left side of each panel, or to release labeled inorganic phosphate from unreacted substrate (Pi). Lanes marked with a dash (-) indicate no enzyme. Wedges indicate decreasing enzyme concentration in each reaction, with the highest concentration of DdiTLP2 WT, KPND, KPHD, NFNKPHD, and DdiThg1 WT corresponding to 14 µM, and the highest concentration of R187D and DCKR corresponding to 90 µM.

Next, we tested activity with the same variants using *D. discoideum* cytosolic tRNA^His^ (cyt-tRNA^His^), for which a labeled GpGpC fragment corresponds to the product of G-1 addition. As previously shown, wild-type DdiThg1 and DdiTLP2 exhibit reversed activity on this tRNA consistent with their known biological roles, with DdiThg1 using cyt-tRNA^His^ as a robust substrate, while DdiTLP2 exhibits no detectable activity on this tRNA (**Figure 4B**) (Long et al. 2016). There was no detectable acquisition of Thg1-like activity observed for any of these variants with the Thg1 substrate cyt-tRNA^His^, even with the KPND mutant that mimicked this absolutely conserved Thg1-specific motif (**Figure 4B**).

Despite the use of structure to guide the choice of these amino acid replacements, the possibility of significant disruption of the overall structure of DdiTLP2 cannot be excluded. To test this possibility, the affinity of WT DdiTLP2 and selected variants for mt-tRNA^His^ substrate was measured using electrophoretic mobility shift assays (EMSA) (**Figure 5A**). A predominant band corresponding to DdiTLP2-bound tRNA was visualized in an enzyme concentration-dependent manner and the fraction of total bound tRNA was quantified for wild-type DdiTLP2 and fit to a binding isotherm, yielding K_D,app_ of 1.1 ± 0.1 μM and a Hill coefficient of ∼3, indicative of cooperative binding, as observed previously for other Thg1 family members (**Figure 5B** and **Table 1**). Binding by the KPND and R187D variants, as representatives of the two tested regions, was quantified using the same approach. For the variant with the Thg1-like KPND sequence, the overall binding behavior (K_D,app_ and apparent cooperativity) was nearly identical to that of wild-type DdiTLP2. These data suggest that changes in the overall protein structure and ability to interact with mt-tRNA^His^ substrate are not the driver of the complete loss of catalytic activity with alterations in the “KPXD”-loop. However, despite the R187D variant only comprising a single amino acid change in a very well-predicted part of the enzyme structure, tRNA binding was significantly altered for this variant, which exhibited an ∼15-fold loss in affinity and loss of cooperative binding. Interestingly, R187 or its analog D150 is not predicted to make any contacts to tRNA in the modeled structures (or in either bona fide structure of an enzyme-tRNA complex), but instead is located near the GTP nucleotide that is used for nucleotidyl transfer (step 2) of the reaction (**Figure 3**). Thus, the basis for this residue’s observed impact on tRNA affinity is not clear.

**Figure 5.**
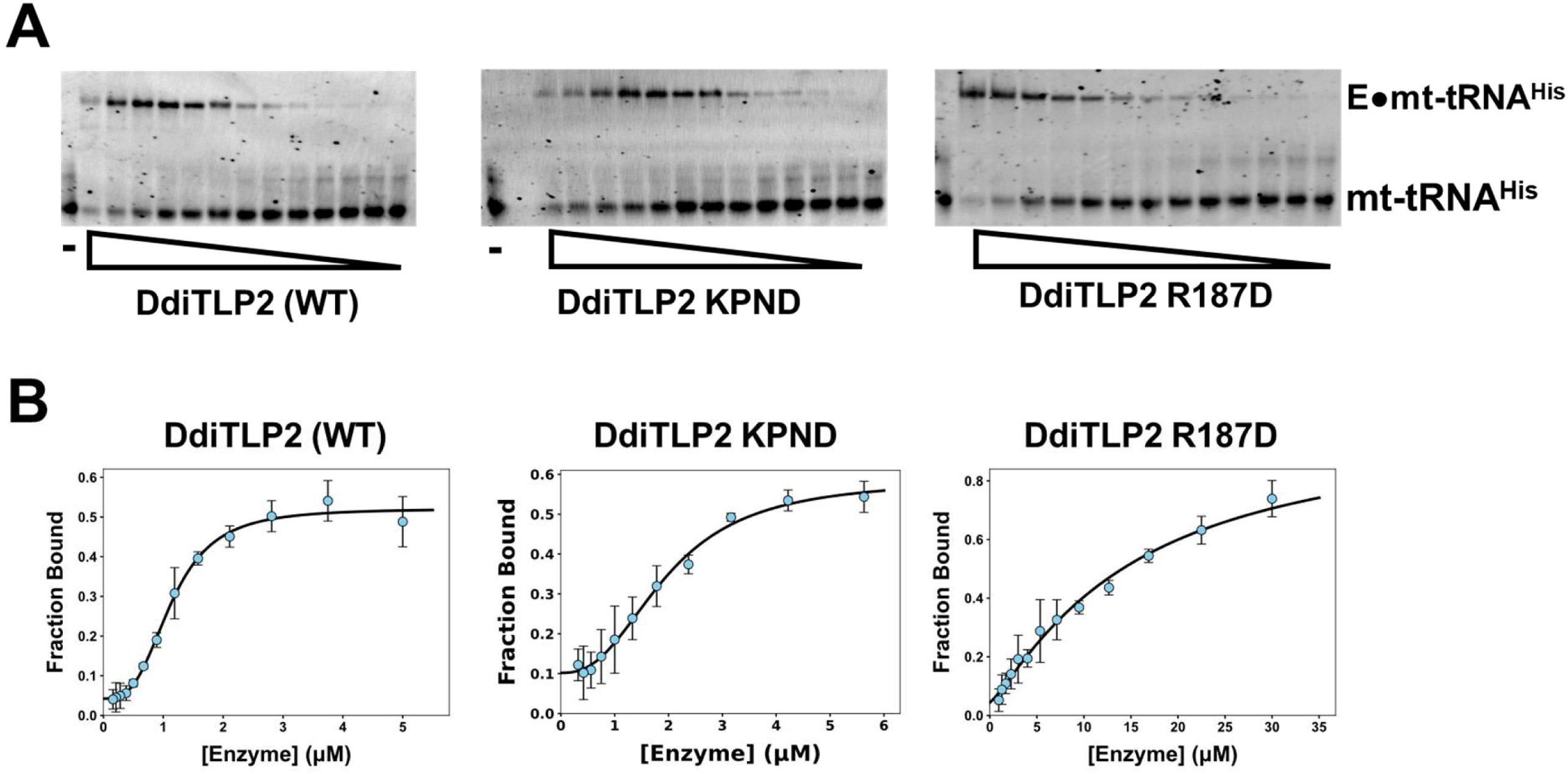
Alterations to unique DdiTLP2 residues affect tRNA binding differently. (A) Binding of DdiTLP2 and variants to mt-tRNA^His^ measured using native PAGE. Representative Electrophoretic Mobility Shift Assays (EMSA) showing the shift of the free mt-tRNA^His^ into higher migrating complexes are shown for WT, KPND and R187D variants, which indicates binding in a concentration-dependent manner, as indicated to the right of each band. Lane marked with – indicates no added DdiTLP2. Binding reactions were assembled with decreasing concentration of excess DdiTLP2 (WT, 0.16 – 5.0 μM; KPND, 0.32 – 5.6 μM; R187D, 0.95 – 30 μM) over free mt-tRNA^His^, and images were obtained using SYBR Gold staining. **(B) Determination of binding parameters from EMSA.** The fraction of bound tRNA was measured at each concentration and fit to eq. 1 to yield K_D,app_ and nH for each binding reaction, as shown in Table 1. Reactions were performed in triplicate, with the average fraction bound at each concentration calculated and plotted with error bars indicating standard error of each measurement.

**Table 1:** Binding of DdiTLP2 variants to mt-tRNA^His^.

| **Table 1: Binding of DdiTLP2 variants to mt-tRNA<sup>His</sup>** |  |  |
| --- | --- | --- |
| **DdiTLP2 enzyme** | **K<sub>D,app</sub><sup>a</sup>**<br>**(μM)** | **nH** |
| Wild-type | 1.1 ± 0.1 | 3.1 ± 0.3 |
| KPND | 2.0 ± 0.1 | 2.5 ± 0.4 |
| R187D | 15.2 ± 5.9 | 1.2 ± 0.3 |

<sup>a</sup> EMSA were performed in triplicate and data were fit to eq. 1 to determine $K_{D,app}$ and nH.

### The single D150R change in DdiThg1 confers TLP-like properties on the enzyme

Because of this result suggesting an unusual role of R187 in DdiTLP2, we created the reciprocal D150R variant in DdiThg1. DdiThg1 D150R also expressed and purified well (**Figure S2A**), but here *in vitro* assays revealed a striking result. While wild-type DdiThg1 exhibits barely any detectable addition of G-1 to mt-tRNA^His^, the “DdiTLP2-like” DdiThg1 D150R variant acquired a dramatically increased activity, producing a similar amount of G-1 addition product (GpGpU) as wild-type DdiTLP2 (**Figure 6A**). Acquisition of DdiTLP2-like activity (WC base pair-dependent G-1 addition across from C73) on mt-tRNA^His^ caused by the single amino acid D150 to R substitution is consistent with the essential nature of an arginine residue at this location for DdiTLP2’s unique ability to add G-1 to this specific tRNA (**Figure 1C**).

**Figure 6.**
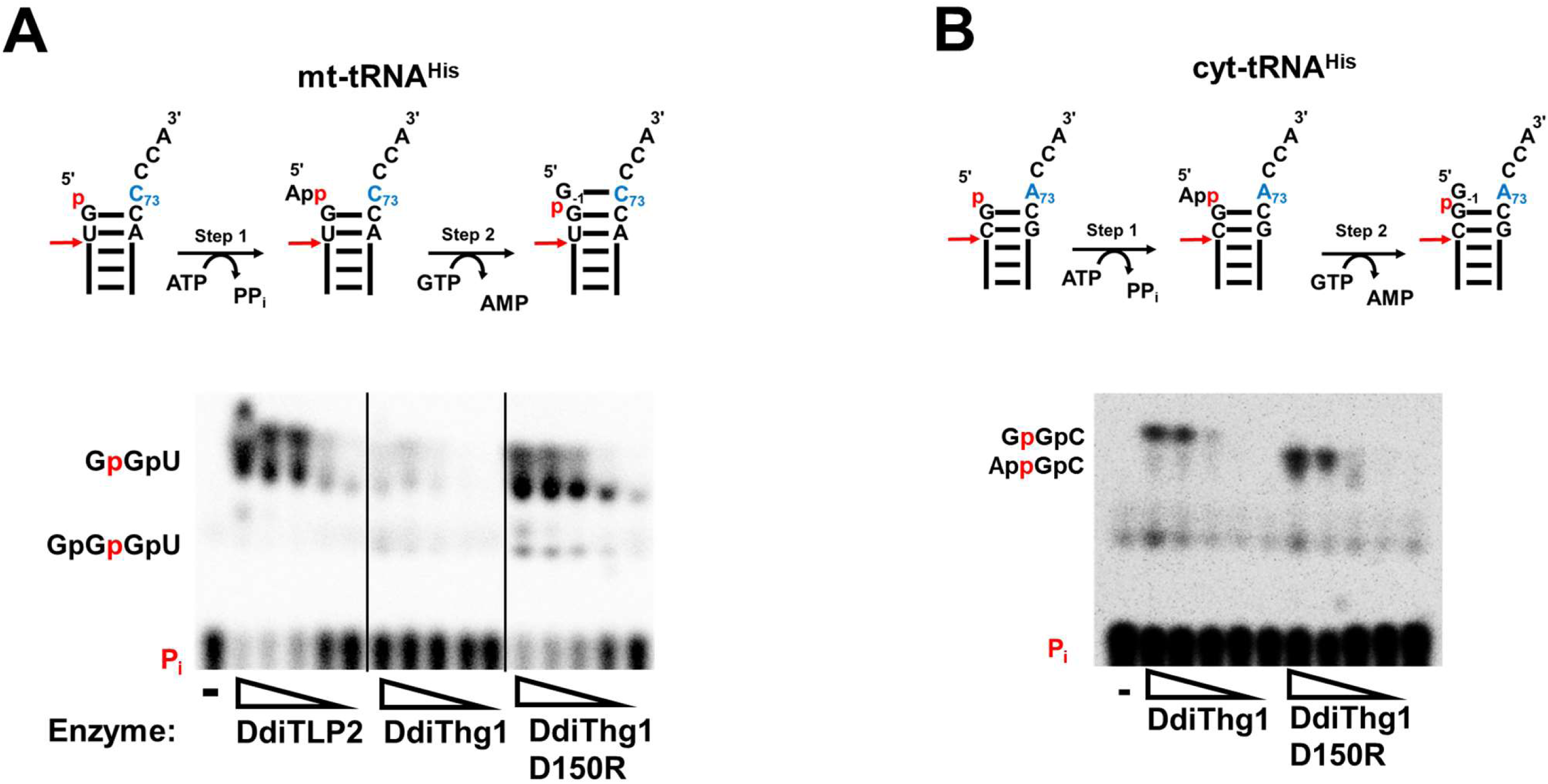
The DdiThg1 D150R variant acquires TLP-like properties. Phosphatase protection assays were performed with 5’-^32^P-labeled transcripts corresponding to **(A) Ddi mt-tRNA^His^** and **(B) Ddi cyt-tRNA^His^.** The indicated enzymes were incubated with labeled tRNA, 0.1 mM ATP and 1 mM GTP for 2 hr, then treated with EDTA to quench the reaction. Reactions were first treated with RNase A to produce shorter labeled fragments, as in the diagrams above each assay with the site of RNase A cleavage shown as a red arrow. Subsequent treatment with phosphatase results in production of the indicated oligonucleotide fragments, or inorganic phosphate (P_i_) from unreacted substrate, which are resolved by TLC. Wedges indicate 5-fold serial dilutions of enzyme, starting from 50 µM. Lanes marked with a dash (-) indicate no enzyme. For mt-tRNA^His^ (**Panel A**), GpGpU is the product of G-1 addition and GpGpGpU is the product of a 2^nd^ (G-2) nucleotide being added, due to the presence of consecutive templating C-nucleotides to act as a template on the 3’-end of this tRNA. The GpGpU spot is observed to smear at high concentrations of enzyme in this solvent system. For cyt-tRNA^His^ (**Panel B**), GpGpC corresponds to the product of G-1 addition, and AppGpC is the 5’-adenylylated intermediate.

We then tested whether D150R alteration impacts DdiThg1 activity on its own physiological substrate, cyt-tRNA^His^ (**Figure 1A**). The D150R DdiThg1 variant retained the ability to recognize and act on cyt-tRNA^His^, based on similar amounts of total products formed at the same enzyme concentrations for variant and wild-type enzymes **(Figure 4B)**. However, the D150R variant produced a completely different type of product than the wild-type enzyme, creating only the lower migrating species (AppGpC) that corresponds to the 5’-adenylated intermediate for this tRNA. Thus, D150R DdiThg1 has a severely impaired ability to proceed after adenylylation through Step 2 to incorporate the non-WC base paired G-1 across from A73 in this cyt-tRNA^His^ (see **Figure 1D**). The strong preference for template-dependent 3’-5’ nucleotide addition activity over the ability to add a non-templated G-1 is a property shared by all TLPs and that distinguishes these enzymes from Thg1-type orthologs (Abad et al. 2010). Thus, this single D to R amino acid change converted the DdiThg1 enzyme into an enzyme that has properties that look more like a typical TLP. These data position the D residue at the heart of the 3’-5’ polymerase activity catalyzed by an ancestral 3’-5’ polymerase, and suggest a possible pathway to acquisition of extant Thg1-type activity by exploiting this conserved amino acid, as explored further below.

### The carboxylate side chain of D150 acts as a gatekeeper for catalysis in DdiThg1

To understand how the D to R change caused this dramatic difference in DdiThg1 substrate specificity and to gain insight into the molecular role of D150, we tested other alterations at this position. Since aspartate and arginine are drastically different in charge, size, and chemical composition, we made replacements to different amino acids chosen to test different aspects of the side chain’s properties. Variants in which DdiThg1 D150 was altered to A, K, L and M were created, thus removing the side chain nearly entirely (A), replacing it with another basic, although smaller, side chain (K), testing a non-polar side chain of similar bulk (L) and altering to a non-polar residue of similar bulk but that also is capable of H-bonding (M). The purified variants were first tested for their ability to add a WC base-paired G-1 to mt-tRNA^His^ (**Figure 7A**). Interestingly, regardless of the side chain replacement made for D150, activity with mt-tRNA^His^ increased substantially relative to the original wild-type DdiThg1. These data indicate that the carboxylate side chain of the wild-type D150 exerts an overall negative effect on catalysis, such that when the D residue is replaced by any other amino acid, the ability to progress through the reaction and incorporate the WC-base paired GTP at position G-1 increases. Thus, in the context of Thg1, D150 acts as a gatekeeper residue to prevent incorrect activity on a non-substrate for this enzyme, such as mt-tRNA^His^.

**Figure 7.**
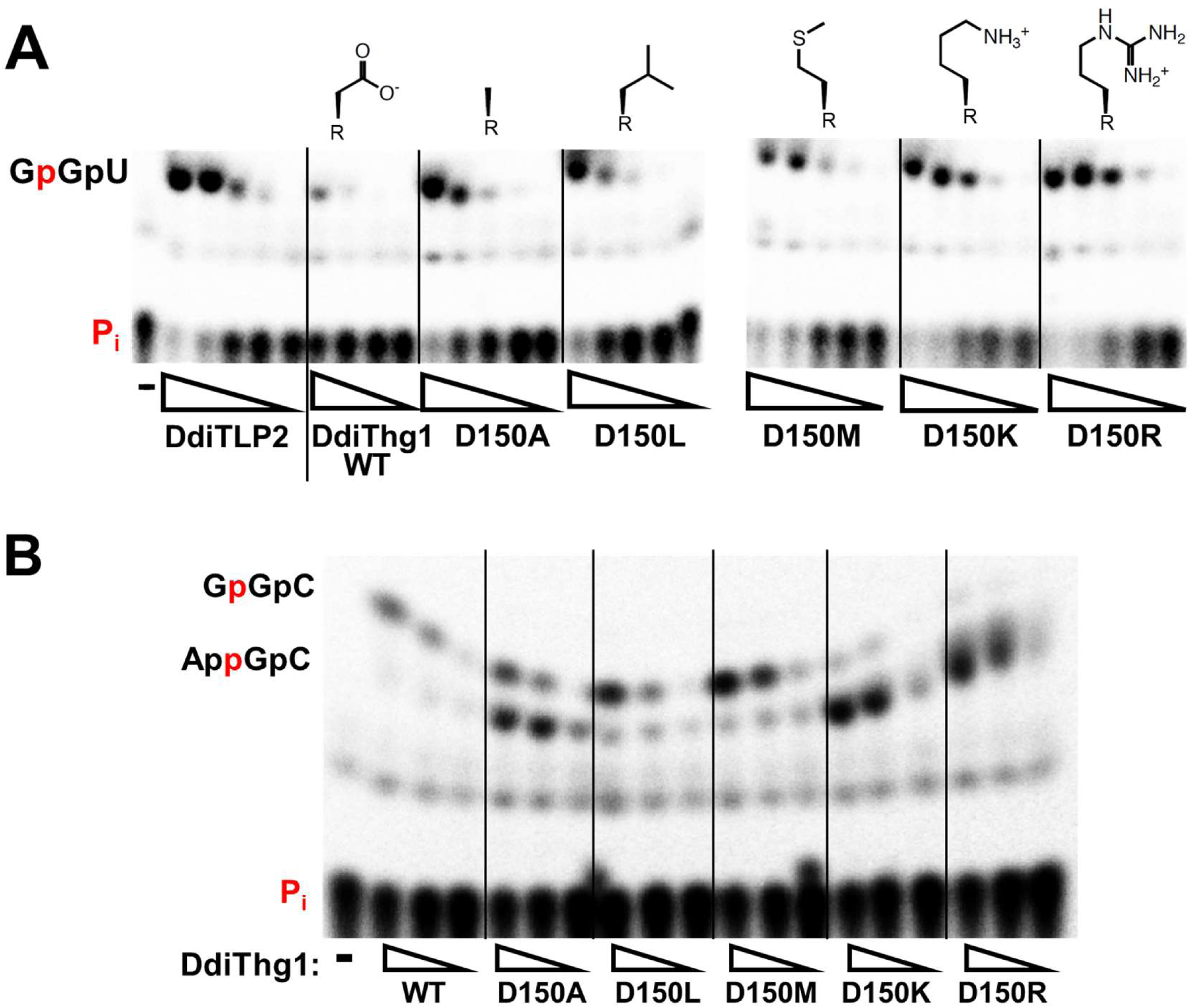
Alteration of D150 to other amino acids has distinct effects on non-WC and WC-templated G-1 addition. Phosphatase protection assays were performed with DdiTLP2 variant enzymes with either **(A) mt-tRNA^His^** or **(B) cyt-tRNA^His^,** with products generated as described in Figure 6. Reactions were initiated by addition of each enzyme (five-fold serial dilutions indicated by wedges, starting at a concentration of 8 μM) to reactions containing 0.1 mM ATP, 1 mM GTP, and 5’-^32^P-labeled tRNA and then were incubated for 2 hr. Lanes marked with a dash (-) indicate no enzyme.

### The identity of the amino acid at position 150 distinctly impacts WC vs. non-WC nucleotidyl transfer steps

We then tested the same D150 variants with the template for non-WC addition of G-1 (Ddi cyt-tRNA^His^). As seen previously with D150R (**Figure 6B**), the D150K variant also caused accumulation of the 5’-adenylated intermediate, suggesting that the presence of a positively charged amino acid at this position allows the enzyme to proceed through Step 1 but does not permit efficient Step 2 nucleotidyl transfer **(Figure 7B)**. However, the D150A variant also had a similar effect, causing accumulation of more 5’-adenylylated tRNA than the wild-type enzyme. Thus, the effect of the D150R alteration is not due solely to reversing the charge at this position. Analysis of the D150L and D150M variants revealed additional complexity in the role of D150 in DdiThg1. These replacements for the carboxylate side chain enabled the enzyme proceeded efficiently through both steps of the reaction, resulting in complete addition of G-1 to cyt-tRNA^His^. Thus, replacement of the bulk of the D150 side chain with similar sized, but uncharged, L or M residues, also restores the enzyme’s ability to also progress through step 2 of the reaction and add GTP to cyt-tRNA^His^.

These results indicate that while any amino acid other than aspartate at the D150 position can support similarly efficient nucleotidyl transfer to mt-tRNA^His^ (because of the presence of similar amounts of G-1 product with all tested variants, **Figure 7A**), efficient nucleotidyl transfer the cyt-tRNA^His^ specifically requires D, L or M at position 150 (**Figure 7B**). Since the main difference between the nucleotidyl transfer steps with mt- vs. cyt-tRNA^His^ is whether the added GTP nucleotide can form a WC base pair (G-1:C73 for mt-tRNA^His^ vs. G-1:A73 for cyt-tRNA^His^, see **Figure 6**), this predicts that the mechanistic role of the D, L or M side chain is only important when the enzyme is trying to incorporate the non-WC base paired GTP to cyt-tRNA^His^. We sought to quantify this specifically using kinetic assays, testing the prediction that the impact of each different amino acid on the kinetics of the nucleotidyl transfer step would be different for a WC vs. non-WC base paired addition. Recently, we developed a new assay that would allow direct measurement of single-turnover rates of the nucleotidyl transfer step (step 2) (Iwaniec et al. 2025). The new assay was necessary because a key reagent needed for assays that had been used previously (i.e, (Smith and Jackman 2012)) to study nucleotidyl transfer kinetics was no longer commercially available. However, this assay requires use of a tRNA transcript that initiates with an A nucleotide (A+1), which is not the case for either of the tRNA^His^ substrates tested up to this point. To use the new assay, we mutated both cyt-tRNA^His^ and mt-tRNA^His^ to initiate with an A+1:U72 base pair in place of the wild-type G+1-C72 (**Figure S4**). Then, each tRNA was transcribed in the presence of [γ-^32^P]-ATP to produce 5’-γ-32P-triphosphorylated tRNA substrates. Addition of G-1 to each of these tRNAs results in release of labeled pyrophosphate (as PP_i_, see **Figure S4A** for reaction scheme), which can be further hydrolyzed to labeled P_i_, and total products quantified using TLC (**Figure 8A**).

**Figure 8.**
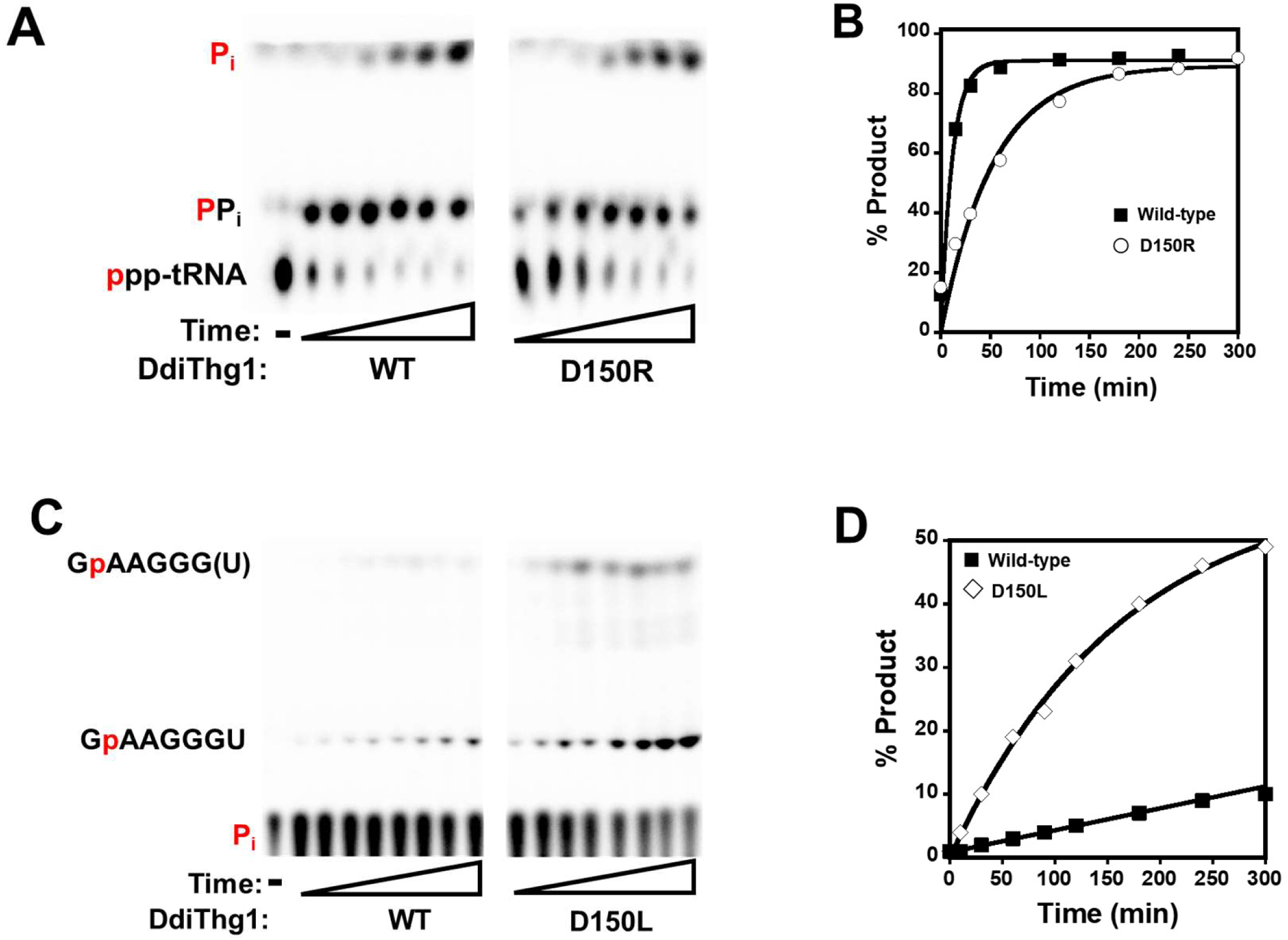
Distinct impact of D150 alterations on single turnover rates of WC vs. non-WC base paired G-1 addition. (A) Representative time courses of nucleotidyl transfer reactions catalyzed by DdiThg1 and D150R variant. To measure rates of nucleotidyl transfer specifically, the indicated enzymes (10 μM) were incubated with 5’-^32^P-labeled triphosphorylated transcripts in which the γ-phosphate has been labeled by ^32^P during *in vitro* transcription. When the enzyme adds a G-1 nucleotide, this causes release of labeled PP_i_ (see Figure 1D), which can be resolved from the unreacted tRNA using TLC. Over time, some PPi hydrolyzes into Pi, which is also quantified as a reaction product. Wedges indicate increasing reaction time from 10-300 min; lane marked with dash (-) correspond to no enzyme control. **(B) Determination of k_obs_ for nucleotidyl transfer for the non-WC base paired addition of G-1 to cyt-tRNA^His^.** Product formation from the time courses shown in (A) was quantified and plotted as a function of time and fit to eq. 2 to determine k_obs_ for each enzyme. Reactions were performed in triplicate and k_obs_ from individual time courses were averaged as reported in Table 2. **(C) Representative time courses used to determine adenylyation rates of DdiThg1 and D150L variant.** Phosphatase protection assays were performed with the indicated enzymes (10 µM) in the presence of 0.1 mM ATP, 1 mM GTP, and 5’-^32^P mt-tRNA^His^. Reactions were quenched by addition of EDTA and treated with RNase A and phosphatase, as described previously. However, the distinct sequence of the mt-tRNA^His^ transcript that was created in order to test nucleotidyl transfer rates results in generation of a distinct product (GpAAGGGU) after G-1 addition. In these assays, a small amount of a higher migrating product that is consistent with further digestion to remove the 3’-U nucleotide was also observed (labeled as GpAAGGG(U) on the TLC image). Both products are clearly enzyme and time-dependent, and therefore were included as products for the quantification of the assays. Wedges indicate increasing reaction time from 15-300 min; lane marked with dash (-) correspond to no enzyme control. **(D) Determination of k_obs_ for adenylylation for the WC base-paired addition of G-1 to mt-tRNA^His^.** Product formation from the time courses shown in (C) was quantified and plotted as a function of time. The reaction with D150L was fit to eq. 2 to determine k_obs_. The reaction with WT DdiTLP2 was analyzed using the method of linear initial rates (see Methods) using eq. 3, based on the linear fit to the data up to 120 min of reaction. Reactions were performed in triplicate and k_obs_ from individual time courses were averaged as reported in Table 2.

**Table 2:** Single turnover reaction rates (k_obs_) for selected DdiThg1 activities.

|  |  |  |  |
| --- | --- | --- | --- |
| **DdiThg1 enzyme** | **k<sub>obs</sub>**<br>**(nucleotidyl transfer)**<br>**cyt-tRNA<sup>His</sup> (non-WC)**<br>**(min<sup>-1</sup>)** | **k<sub>obs</sub>**<br>**(nucleotidyl transfer)**<br>**mt-tRNA<sup>His</sup> (WC)**<br>**(min<sup>-1</sup>)** | **k<sub>obs</sub>**<br>**(adenylylation)**<br>**mt-tRNA<sup>His</sup> (WC)**<br>**(min<sup>-1</sup>)** |
| Wild-type | 0.07 ± 0.01 | 0.54 ± 0.02 | 0.0007 ± 0.0004<sup>a</sup> |
| D150A | 0.030 ± 0.006 | 0.66 ± 0.06 | 0.009 ± 0.005 |
| D150L | 0.14 ± 0.01 | 3.0 ± 0.3 | 0.010 ± 0.006 |
| D150M | 0.18 ± 0.07 | 3.4 ± 0.6 | NT<sup>b</sup> |
| D150K | 0.019 ± 0.002 | 0.8 ± 0.2 | NT<sup>b</sup> |
| D150R | 0.017 ± 0.002 | 0.9 ± 0.1 | 0.03 ± 0.01 |

<sup>a</sup> k<sub>obs</sub> was estimated using the method of linear initial rates
<sup>b</sup> Not tested.

Using this assay, time courses were performed under single-turnover ([E] >> [S]) conditions at saturating (10 μM) enzyme concentrations and product formation was fit to the single-exponential rate equation to derive k_obs_ for the nucleotidyl transfer (step 2) of the non-WC vs. WC G-1 addition reactions (on cyt-tRNA^His^ and mt-tRNA^His^, respectively) (**Figure 8B**, **Table 2**). The effects on the non-WC nucleotidyl transfer step with cyt-tRNA^His^ showed the expected pattern, with the D150A, K and R variants that accumulated the step 1 adenylylated intermediate correspondingly exhibiting slower (∼2-4-fold decreased) k_obs_ values than wild-type DdiThg1 (**Figure 8B**, **Table 2**). The D150L and D150M variants showed modestly increased k_obs_ for non-WC addition to cyt-tRNA^His^, confirming that the presence of the non-polar side chain is indeed able to support robust nucleotidyl transfer activity as inferred from the complete assay shown in **Figure 7B**. We used the same assay to similar measured single turnover rates for WC base-paired nucleotidyl transfer with mt-tRNA^His^. For this WC base-paired nucleotidyl transfer reaction, A, K and R variants exhibited k_obs_ values that were similar to, or even slightly increased compared to wild-type (**Table 2**). Again, the rates observed with the L and M substitutions were also substantially increased compared to wild-type DdiThg1. Even though there is clearly a positive impact of L or M at position 150 on the actual rate of step 2 (nucleotidyl transfer) regardless of the identity of the base pair being formed, this does not lead to an increase in the overall G-1 addition product for these variants relative to others with slower nucleotidyl transfer rates in assays that require both step 1 and step 2 (**Figure 7**) because of the rate-determining nature of the first, 5’-adenylylation step, as demonstrated previously (Smith and Jackman 2012).

### The D150 residue plays a dual role in Thg1-type enzymes that may have been exploited during evolution of eukaryotic G-1 addition activity on tRNA^His^

Taken together, these results show that in the context of DdiThg1, D150 plays two distinct roles. D150 acts as a negative element restricting overall enzymatic activity in the context of its non-substrate, mt-tRNA^His^ (**Figure 7A**). The distinct antideterminants in *D. discoideum* mt-tRNA^His^ that prevent its recognition for G-1 addition by DdiThg1 despite the fact that it contains the otherwise necessary GUG anticodon are not yet known. However, this negative role for D150 specifically requires the presence of the carboxylate side chain, since replacement with any other tested amino acid allows DdiThg1 activity on this non-cognate tRNA (**Figure 7A**). D150 also acts separately as a positive element promoting efficient nucleotidyl transfer of a non-WC base paired GTP onto its cognate cyt-tRNA^His^ substrate, since removing the side chain or replacing it with a positively charged residue results in decreased rates of non-WC nucleotidyl transfer (**Figure 7B**, **Table 2**). For this second role, however, the charged carboxylate side chain is not essential, since it can also be replaced with similarly sized non-polar side chains (L or M) and still exert the same positive effect (**Figure 8**, **Table 2**). Thus, different chemical characteristics of the same amino acid are apparently responsible for each of these effects. So, how can these disparate effects on catalysis be explained?

The modeled structures of the DdiThg1 vs. DdiTLP2 enzymes generated here combined with recent biochemical insight into the nature of the two nucleotide binding sites in the enzyme provide a possible molecular basis for this observed dual role for D150. The models were generated with all three substrates that are utilized by Thg1/TLP during the two-step reaction mechanism: tRNA^His^, ATP for adenylylation and GTP for nucleotidyl transfer **(Figure S2 and Figure 3).** The choice to simultaneously include all three substrates in each structure, despite the fact that the ATP and GTP nucleotides are predictably involved in separate sequential reaction steps (**Figure 1D**), is based on our recent demonstration that both nucleotides are already bound at the active site of a bacterial TLP prior to the rate-determining adenylylation step (Iwaniec et al. 2025). In these previous studies, we demonstrated that the TLP uses information about whether the correct WC base paired NTP is bound at the nucleotidyl transfer site to dictate whether the rate-determining adenylylation step can proceed, and in the absence of a correct WC base-pairing NTP that matches the templating nucleotide, the rate of 5’-adenylylation is nearly undetectable (Iwaniec et al. 2025). Interestingly, in the structural model of DdiThg1 in the presence of tRNA, ATP and GTP (**Figure 3**), we observed a potential hydrogen-bonding interaction between the carboxylate side chain of D150 and the 3’-OH of the GTP bound in the nucleotidyl transfer site. This contact had never been previously observed in the only other available Thg1-tRNA complex structure because this structure lacked NTPs and also had poor resolution of tRNA nucleotides in the active site (Nakamura et al. 2013). Importantly, this GTP 3’-OH will become the nucleophile to attack the product of step 1 (adenylylation) during step 2 of the reaction, but predictably can not enter a catalytically productive conformation where it would be coordinated to the catalytic metal ions until step 1 is completed. The nature of the conformational change that allows the transition of substrates and intermediates to occupy different positions relative to the catalytic metal ions between step 1 and step 2 is unknown.

Thus, the negative effect of D150 on the rate-determining adenylylation step may be related to its ability to use the carboxylate-GTP interaction to hold the incoming NTP in this catalytically unproductive conformation until the correct features of a particular tRNA-NTP combination are determined, at which point the adenylylation step can proceed. The catalytic features that would trigger “release” of the GTP 3’-OH nucleophile to the active conformation remain to be determined, and identification of these will require additional structural and mechanistic characterization.

Directly measuring the predicted effect of D150 on the 5’-adenylylation step is difficult because, as we described above, the maximal rate of adenylylation can only be achieved in the presence of the correct nucleotidyl transfer NTP, and thus isolated measurements of step 1 rates in the absence of the next incoming NTP likely underestimate the actual rate of this step and may impact the ability to test the roles of key residues. Here, we took advantage of the fact that 5’-adenylylation is clearly rate-limiting for DdiThg1 as described above, and used the phosphatase protection assay to measure single turnover rates of 5’-adenylylation with wild-type Thg1 and D150A, D150L and D150R variants by measuring activity on the non-cognate mt-tRNA^His^ in the presence of both ATP and GTP (**Figure S4B**). Total G-1 product formation was quantified as a function of time to determine an overall k_obs_ for 5’-adenylylation based on the fact that adenylylation is the first and rate-determining step for this reaction **(Figure 8C and 8D).**

The measured rates agree with the predicted role of D150 as a negative gatekeeper residue for 5’-adenylylation, since replacement of the inhibitory carboxylate side chain with any of these different amino acids results in a significantly increased rate for the adenylylation step of 10-40-fold compared to wild-type when the enzyme is adding a WC base paired NTP, such as with mt-tRNA^His^ **(Figure 8D**, **Table 2).**

The molecular basis for the second role of D150 in promoting the rate of the nucleotidyl transfer step for a non-WC base paired addition of G-1 is less clear. However, this observation may provide an explanation for the evolution of eukaryotic Thg1 enzymes’ ability to recognize two different kinds of base pairs (WC and non-WC) using the same active site. In this scenario, the ancestral Thg1/TLP would already have developed a mechanism that depends on utilizing the interaction between D150 and the incoming GTP to improve fidelity of WC base pair formation, as described above. Possibly, the precursor to eukaryotic Thg1 took advantage of the D150 residue’s ability to position incoming NTPs in the active site to also facilitate the positioning of a GTP nucleotide that would otherwise not be stably bound in the active site because it is not forming a WC base pair. For this second role of stabilizing a non-WC bound GTP to promote this nucleotidyl transfer step (**Table 2**), the important properties of the D residue only require steric effects that can also be achieved with L or M at this position. Nonetheless, changes at D150 would not be allowed because they would impact the primary gatekeeper function of this residue. It is interesting to speculate whether DdiTLP2 may have been required to alter this otherwise universally conserved residue to acquire its ability to catalyze specific incorporation of the G-1 into mt-tRNAHis in an orthogonal reaction to that of DdiThg1, and thus preserve the fidelity of tRNA^His^ aminoacylation in *D. discoideum* mitochondria.

These results may also provide an explanation for the puzzling phenotype that was originally attributed to the D153A alteration in ScThg1. Consistent with the observations here with DdiThg1, the ScThg1 D153A variant also showed increased G-1 addition activity (3-fold greater than wild-type ScThg1) in the *in vitro* assays (Jackman and Phizicky 2008). Yet, the observed growth defect when this variant was used as the only source of Thg1 activity in *S. cerevisiae* seemed at odds with the fact that this variant was more active at the biologically-required G-1 addition reaction. It is possible that, like with DdiThg1, alteration of this residue eliminates the gatekeeper function that enhances Thg1 fidelity, and therefore expressing this enzyme *in vivo* leads to inappropriate G-1 addition to other incorrect (non-His) tRNA(s), with a corresponding negative effect on growth. Thus, it is possible that the universally conserved nature of the D at this position in all Thg1/TLP enzymes is the result of a tradeoff between efficiency that could be achieved with other substitutions at this position but would come at a cost of reduced overall fidelity and specificity.

## CONCLUSIONS

Here, we used sequence comparison of an unusual member of the Thg1/TLP 3’-5’ polymerase family (DdiTLP2) to identify sequences that are the source of this enzyme’s unusual specificity for one substrate, mitochondrial tRNA^His^. Residues in a conserved loop region that surround the binding site for an ATP used in 5’-adenylylation appear to facilitate the unique biochemical properties of DdiTLP2, since changes to these amino acids completely disrupted catalysis without affecting apparent interaction with the tRNA (**Figures 3, 4 and 5**, **Table 1**). However, this also led us to uncover two distinct biochemical functions for another universally conserved residue (D150) in the fidelity of 3’-5’ polymerase activity (**Figures 6, 7 and 8**, **Table 2**). Based on these results we propose that the D150 residue played a critical role in the evolution of the non-WC base paired addition of G-1 that is the essential activity of extant eukaryotic Thg1 enzymes. These results also suggest a structure and mechanism-based explanation for a previously observed connection between the two steps of Thg1/TLP-mediated catalysis, in which the presence of the correct NTP for step 2 is able to exert an effect on the first, rate-determining step (Matlock et al. 2019; Patel et al. 2021; Iwaniec et al. 2025). Future structural and biochemical studies will be required to provide explicit evidence for this proposed evolutionary pathway and the specific gatekeeper function of D150, but the picture of 3’-5’ addition catalyzed by these non-canonical polymerases is becoming increasingly clear through studies of the unique enzymes of the *D. discoideum* system.

## MATERIALS AND METHODS

### Sequence alignment

Pairwise sequence alignment was performed by EMBOSS Needle (Madeira et al. 2024). Multiple sequence alignment was performed in CLUSTAL O (1.2.4) with 12 Thg1/TLP enzymes from all three domains of life: *Dictyostelium discoideum* (Ddi) Thg1, TLP2, TLP3, and TLP4; *Bacillus thuringiensis* (Bt) TLP; *Acanthamoeba castellanii* (Aca) TLP1 and TLP2; *Homo sapiens* (Hs) Thg1; *Saccharomyces cerevisiae* (Sc) Thg1; *Candida albicans* (Ca) Thg1; *Methanosarcina acetivorans* (Ma) TLP; and *Methanopyrus kandleri* (Mk) TLP (Sievers and Higgins 2014).

### AlphaFold 3 Modeling

Each enzyme was modeled using AlphaFold 3 server (Abramson et al. 2024). Each protein was modeled as a homotetramer with two substrate tRNA, twelve Mg ions, four ATP, and four GTP. pTM was above 0.5 for all structures. ipTM was 0.43 for DdiTLP4, 0.68 for DdiTLP2, and 0.87 for DdiThg1. Structures were visualized using the PyMOL Molecular Graphics System, Version 2.4.1 Schrödinger, LLC.

### Thg1/TLP protein expression and purification

Plasmids for expression and purification of ScThg1, DdiThg1, and DdiTLP2 enzymes with N-terminal His6-tag sequences have been described previously.(Abad et al. 2011; Iwaniec et al. 2025) Variants of these enzymes were made using the Phusion site-directed mutagenesis kit (Thermo Scientific). Briefly, enzymes were expressed in *E. coli* BL21 (DE3) pLysS or Rosetta pLysS cells and overexpressed proteins were purified using immobilized metal-ion affinity chromatography (IMAC) with TALON resin (Clontech).(Abad, Rao, and Jackman 2010) Proteins were dialyzed after elution from the column in a dialysis buffer containing 20 mM Tris pH 7.5, 500 mM NaCl, 4 mM MgCl_2_, 50% Glycerol, 1 μM ethylenediaminetetraacetic acid (EDTA) pH 8.0 and 1 mM dithiothreitol (DTT) and stored at -20°C. The BioRad protein assay was used to determine the concentrations of purified proteins for assays described below. On the day of use in assays, proteins were diluted as needed using enzyme dilution buffer (20 mM Tris pH 7.5, 0.5 mg/mL bovine serum albumin (BSA) (Invitrogen), 500 mM NaCl, 1 mM DTT).

### *In vitro* transcription of tRNAs

T7 RNA polymerase was used for runoff *in vitro* transcription using digested plasmids encoding tRNA^His^ wild-type and variant constructs downstream from the T7 RNA polymerase class III consensus Φ6.5 promoter, as previously described (Iwaniec et al. 2025). tRNA variants to introduce A+1-U72 base pairs for nucleotidyl transfer assays were created using the QuickChange site-directed mutagenesis kit (Agilient). tRNA variants starting with A+1 were *in vitro* transcribed using the P266L mutant of T7 RNA polymerase under the Φ2.5 promoter to improve initiation with the unfavorable A+1 residue (Guillerez et al. 2005).

### Production of 5’-[32P]-monophosphorylated tRNA

5’-[^32^P]-labeled tRNA was generated by *in vitro* transcription followed by calf intestinal phosphatase (NEB) treatment and labeling with T4 polynucleotide kinase (NEB), as previously described (Jackman and Phizicky 2006a; Jackman and Phizicky 2008).

### Phosphatase protection assay for G-1 addition by Thg1/TLP enzymes

G−1 addition reactions were performed at room temperature by reacting 5′-monophosphorylated [^32^P]-tRNA^His^ substrate with 0.1 mM ATP and 1 mM GTP in the presence of Thg1/TLP for two hr. The reaction buffer contained 25 mM HEPES pH 7.5, 10 mM MgCl_2_, 125 mM NaCl, 0.2 mg/mL BSA, and 3 mM DTT. To quench the reactions, EDTA was added to the samples to a final concentration of 50 mM. The samples were then incubated with 1 mg/mL of RNaseA (Ambion) at 50°C for 10 min. RNase A-digested samples were treated with 0.1 U/µL calf intestinal phosphatase (CIP) (Invitrogen) and incubated at 37°C for 30 min. The products were resolved using silica thin-layer chromatography (TLC) in an *n*-propanol:NH_4_OH:H_2_O (55:35:10) solvent system. as described previously (Jackman and Phizicky 2006b). TLC plates were phosphor-imaged using Amersham Typhoon and quantified by the Image Quant program.

### Electrophoretic Mobility Shift Assay (EMSA) for tRNA binding

DdiTLP2 wild-type or variant enzyme was diluted using 20 mM Tris pH 7.5, 500 mM NaCl, 1 mM DTT. Enzyme dilutions and 2 ng/uL Ddi mt-tRNA^His^ were incubated at room temperature for 30 min in 20 mM Tris pH 7.5, 2 mM MgCl_2_, 100 mM NaCl, 2 mM DTT, 0.002% Tween 20. Reactions were resolved on native 6% acrylamide/bis-acrylamide 29:1 (Bio-Rad), 0.5X TBE (44 mM Tris adjusted to pH 8.3 using boric acid, 1.25 mM Na_2_EDTA) 2 mM MgCl_2_ gels. Gels were pre-run in 0.5X TBE at 4°C for one hr at 90V. Loading buffer (final concentration in samples of 2.5% Ficoll 400, 0.16X TBE) was added to RNA/enzyme samples, and then samples were subjected to electrophoresis at 4°C and 90V. Gels were stained with SYBR Gold (Invitrogen) for 15 - 30 min and then rinsed with deionized water twice to remove stain. Gels were then fluorescently imaged using the Cy2 laser (488 nm) on Amersham Typhoon and quantified using Image Quant program. *K_D_* was determined by fitting to the equation below using Python. Eq. 1 was used to calculate *K_D_* from the plot of % bound (F) vs. enzyme concentration [E] where *K_D,app_* is the dissociation constant, nH is the Hill coefficient, and minimum and maximum % bound were also obtained for the fit of each binding isotherm.

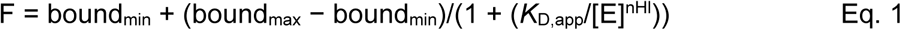

### Single turnover nucleotidyl transfer assays

5’- [γ-^32^P] triphosphorylated-tRNA^His^ A+1-U+72 tRNA^His^ substrates (cyto- and mt-tRNA^His^) were produced via *in vitro* transcription by T7 polymerase P266L mutant in the presence of γ-[^32^P]-ATP (6000 Ci/mmol, Revvity) (**Figure S4A**). Nucleotidyl transfer reactions with [^32^P]PP-tRNA^His^ A1-U72 substrates were performed at 25°C with 1 mM GTP, and initiated by the addition of 10 μM enzyme, as indicated. For reactions with mt-tRNA^His^, the reaction buffer contained 25 mM HEPES pH 7.5, 10 mM MgCl_2_, 125 mM NaCl, 0.2 mg/mL BSA, and 3 mM DTT. For reactions with cyt-tRNA^His^, the reaction buffer contained 25 mM bis-tris pH 6.0, 10 mM MgCl_2_, 50 mM NaCl, 0.2 mg/mL BSA, and 3 mM DTT. At selected timepoints, 5 µL of reaction was quenched with 0.5 µL of 500 mM EDTA and 0.5 µL of 10 mg/mL RNaseA (Ambion) and placed on ice. After the reaction was completed, all quenched samples were incubated at 50°C for 10 min to allow complete digestion of the labeled tRNA to shorter fragments (see **Figure S4B**). Enzymes were precipitated from the RNase A-digested samples by adding 1 µL of trichloroacetic acid (TCA), incubating on ice for 5 min, and spinning down at room temperature for 2 minutes at 15,000 rpm. The products were then resolved using PEI cellulose thin-layer chromatography (TLC) in a 0.5 M KPO_4_, pH 6.3: methanol (80:20) solvent system to separate the labeled pyrophosphate (P*P_i_) from unreacted labeled tRNA, as described previously (Smith and Jackman 2012). TLC plates were phosphor-imaged using Amersham Typhoon and quantified by the Image Quant program. Eq. 2 was used to calculate *k_obs_* using Kaleidagraph (Synergy software), where % product at each time point (%Pt) was plotted vs. time and fit to the single exponential rate equation, where ΔP is maximal product formation, to determine kobs for each reaction. Reactions with each enzyme were performed at least in triplicate, with the reported kobs determined as the average of the three independent trials.

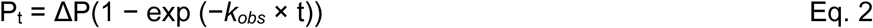

### Single turnover adenylylation assays

Adenylation rates were measured by performing G-1 addition assays on 5’-monophosphorylated [^32^P]-mt-tRNA^His^ substrate at 25°C with 0.1 mM ATP and 1 mM GTP in the presence of a saturating concentration of enzyme (10 µM). Reaction buffer contained 25 mM HEPES pH 7.5, 10 mM MgCl_2_, 125 mM NaCl, 0.2 mg/mL BSA, and 3 mM DTT. At varied timepoints, 5 µL of reaction was quenched with 0.5 µL of 500 mM EDTA and 0.5 µL of 10 mg/mL RNaseA (Ambion) and placed on ice. After the reaction was completed, all quenched samples were incubated at 50°C for 10 min. RNase A-digested samples were treated with 0.1 U/µL CIP)(Invitrogen) or 0.2 U/µL Quick CIP (New England Biolabs) and incubated at 37°C for 30 min. The products were resolved using silica thin-layer chromatography (TLC) in an *n*-propanol:NH_4_OH:H_2_O (55:35:10) solvent system as described previously (Jackman and Phizicky 2006b; Smith and Jackman 2012). TLC plates were phosphor-imaged using Amersham Typhoon and quantified by the Image Quant program. Since adenylylation is the first reaction step and reactions were performed under single turnover conditions (enzyme in excess of substrate), % adenylylation was calculated by quantifying all product spots divided by the reactant inorganic phosphate spot in each lane. Single turnover k_obs_ were determined for each time course using Eq. 2, and rates were measured in at least three independent assays, which were averaged for the reported k_obs_.

For wild type DdiThg1, very slow adenylylation was observed (< 25% conversion to products after 5 hours of reaction). Therefore, k_obs_ was estimated using the method of initial rates, according to Eq.3, where v_o_ is the linear initial rate derived from the slope of the fraction product vs. time plots and ΔP_t_ is the maximal fraction of product conversion determined separately using DdiTLP2 and the mt-tRNA^His^ substrate.

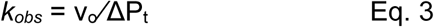

## ACKNOWLEDGMENTS

The authors would like to acknowledge NIH R01 GM087543 and R01 CA260414 (to J.E.J) and NIH T32 GM141955 (to G.N.J.) for funding support.

## Notes

### Competing Interest Statement

The authors have declared no competing interest.

